# Predicting Influenza Virus Host Tropism and Zoonotic Spillover Risk from Protein Sequences

**DOI:** 10.64898/2026.05.21.726772

**Authors:** Alexandra K. Longest, Taylor M. Grace, Minh Tran, Blake Northrop, Ashlyn Donohue, Ahmad Said, Stephanie L. Guertin, Brian E. Root

## Abstract

Novel infectious diseases, predominately originating from non-human animals, pose a significant threat to global public health and economic stability. Avian influenza virus presents an especially significant challenge due to its high mortality rates and spillover capability into new host species. Recent H5N1 spillover events into poultry and cattle resulted in massive economic burden and increased human health risk. Traditional methods of disease surveillance rely on reactive case detection and pathogen characterization, providing insufficient lead time for effective intervention. Computational tools that allow efficient and proactive prediction of zoonotic potential are critical in mitigation of influenza outbreaks and identification of strains with human spillover risk. Existing models predicting influenza virus subtypes or host have been developed; however, the complexity of spillover events, including the non-binary nature of zoonotic potential, limits the capabilities of these models. In the approach reported here, rich protein language model embeddings were generated from ESM-2 for each protein in influenza virus strains and used to predict the protein host tropism probabilities across nine animal families. The protein host tropism model achieved weighted precision and recall scores of 0.95 and 0.95, respectively. We then constructed a zoonotic risk prediction model using the outputs from the protein host tropism prediction model to classify the strains into six classifications: avian, mammal, human, avian-to-human zoonotic, avian-to-mammal zoonotic, or mammal-to-human zoonotic. The average weighted precision and recall scores for this model were 0.90 and 0.90, respectively. This framework advances the prediction of influenza zoonotic risk by being agnostic to influenza subtype, incorporating non-human mammals and mammal zoonotic spillover classifications, and using the full influenza proteome to capture the complexity of spillover dynamics.

## Introduction

The emergence of novel infectious diseases, most of which originate from non-human animals, significantly threatens global public health and economic stability [1]. The COVID-19 pandemic heightened awareness that such future epidemic or pandemic threats are inevitable, with zoonotic viruses representing a particular concern. Among these zoonotic threats, influenza viruses warrant special attention due to their high mutation rates, capacity for interspecies transmission, and history of causing global pandemics [2]. Avian influenza viruses (AIV) currently inflict substantial economic losses on the US and global agricultural sectors [3]; however, their significance extends beyond economic impact. When AIV spillover into humans, they pose a serious threat to public health. The immediate and lasting impacts of spillover events from the last 100 years underscore the urgent need for improved strategies to identify and assess the pandemic potential of AIV before they cross the species barrier [4–6].

While the need to monitor, assess, and predict this threat is clear, doing so has proven extremely challenging. Traditional methods of disease surveillance rely on the detection of animal and human cases and subsequent pathogen characterization. To counter the threat of zoonotic spillover, several strategies have been deployed, including genomic surveillance and virus discovery in animal host reservoirs [4,6,7]. Frameworks such as the World Health Organization’s (WHO) Tool for Influenza Pandemic Risk Assessment (TIPRA) use a systematic process for assigning level of risk [8]. The hazard assessment includes virological, epidemiological, and clinical information to take a comprehensive approach to the evaluation. However, some of these criteria, such as lab animal transmission models and receptor binding properties, are time and resource intensive [5], often resulting in a reactive approach and insufficient lead time for effective intervention.

Recent advances in modeling and computational analysis offer new opportunities for predicting spillover events. These methods often utilize host-associated genetic markers and phylogenetic analysis to identify strains with zoonotic potential [9]. However, mutations and reassortments identified in past zoonotic events may not be detected in other spillover events [9–11]. Models predicting zoonotic potential based on viral genome can identify sequences associated with current, circulating human-infecting strains [12], but these approaches risk deprioritizing identification of generalizable signals of human infection [5], resulting in poor prediction of novel AIV strains. A computational tool that allows for efficient and effective prediction of zoonotic potential is therefore essential for mitigating the risk of future influenza outbreaks.

Models that predict the AIV subtype or host tropism based on genomic and protein sequence have been developed. Park *et al.* [13] developed the PAIVS (Prediction of Avian Influenza Virus Subtype) tool to predict AIV subtype, including data pre-processing, genomic assembly, variant calling, and interpretation of next-generation sequencing data. Flu-level Convolutional Neural Network (Flu-CNN) [14] and WaveSeekerNet [15] use different machine learning (ML) approaches to assign the influenza subtype and host source. These models flag potential cross-species transmission events in cases where the host assigned by the model does not match the host from which the sample was obtained. However, Flu-CNN only focuses on transmission to humans, and WaveSeekerNet focuses on host prediction classification rather than incorporating a zoonotic spillover risk category [14,15]. These models reflect zoonotic potential as binary, but infection by strains less fit for crossover can occur when exposed to higher doses of the virus [16].

Eng *et al.* [17] developed a model for predicting avian-to-human zoonotic risk by treating zoonotic strains as their own classification, but this model did not include non-human mammals. Non-human mammals, such as cattle and pigs, have major economic impact on agricultural industries, and can also be intermediate reservoirs for AIV [18]. In these hosts, AIVs could mutate or undergo coinfection with another influenza virus, which could result in reassortment and the emergence of a new strain capable of crossing additional species barriers [2]. These events increase the risk that new variants emerge, infect humans, and potentially develop transmission capability between humans. Pigs are of high concern because they possess both avian and human receptor binding sites, which increases viral reassortment potential [19]. This property has led to their designation as “mixing vessels.” A spillover from pigs caused the 2009 swine flu pandemic [20]. Therefore, a zoonotic risk prediction model that includes non-human mammals is crucial for AIV pandemic prevention. Such a model could guide experts on where to allocate additional resources for biosurveillance and laboratory work and provide early warnings for strains of concern.

We applied a systems approach in which protein host tropism was first characterized then ensembled to predict zoonotic risk, complementing conventional techniques that investigate amino acid positions that show strong selection for specific hosts by identifying zoonotic potential represented in the variability of mutation and reassortments seen in documented spillover events. We input rich protein language model embeddings of each individual protein in an influenza strain in a ML model to predict the protein host tropism probabilities to each of nine animal families. In this first model, individual protein sequences commonly detected in more than one family may be represented as equal- or near-equal probabilities between those families. These familial probabilities were used as input features to train a second ML model on zoonotic influenza A viruses (IAV). The second model generates probabilistic classification outputs, quantifying the likelihood of each strain belonging to either a host family group or zoonotic spillover category. These values were designated as the risk prediction value, with the highest value determining the strain classification.

## Materials and methods

### Data collection and preparation

Protein sequences were obtained from the National Center for Biotechnology Information (NCBI) (https://www.ncbi.nlm.nih.gov/protein) and Global Initiative on Sharing All Influenza Data (GISAID) (https://gisaid.org/) databases on October 29^th^, 2024 and November 18^th^, 2024, respectively. All sequences were analyzed and processed to obtain the highest quality data for training the model. Protein sequences deviating by more than ±10% in length from the reference strain (canonical influenza A sequence *A/Puerto Rico/8/1934* H1N1) were removed from the dataset. Any sequences with non-standard amino acids were removed. Additionally, sequences were required to have the following metadata: accession number, nucleotide, species, genus, family, length, genotype, segment, protein, country of isolate, host, collection date. Duplicates between NCBI and GISAID databases were cross-checked using accession number/isolate name and subsequently removed. Finally, the data was subset to only include nine identified host families of interest, with sequences from others removed. After processing, the complete dataset comprised 2,711,780 unique sequences.

Zoonotic IAVs were identified based on published literature, including reports from the United States’ Center for Disease Control (CDC) and the WHO, to ensure a high quality confirmed zoonotic dataset. This dataset tracked the host species and originating spillover species; using this information, we classified the host family groups (i.e., avian, human, mammal) and zoonotic spillover categories (i.e., avian-to-human, avian-to-mammal, mammal-to-human). The resulting 248 confirmed zoonotic IAVs, excluding those from Eng *et al.* [17], are detailed in Tables 1 and 2.

**Table 1.** Geographic summary of confirmed zoonotic IAVs from literature review.

| Country | H1N1 | H1N2 | H3N2 | H3N8 | H5N1 | H5N2 | H5N5 | H5N6 | H5N8 | H7N9 | H9N2 | H10N5 | H10N8 | Total |
| --- | --- | --- | --- | --- | --- | --- | --- | --- | --- | --- | --- | --- | --- | --- |
| Argentina |  |  |  |  | 2 |  |  |  |  |  |  |  |  | 2 |
| Brazil | 1 | 1 |  |  |  |  |  |  |  |  |  |  |  | 2 |
| Cambodia |  |  |  |  | 11 |  |  |  |  |  |  |  |  | 11 |
| Canada |  |  |  |  | 2 |  | 3 |  |  |  |  |  |  | 5 |
| Chile |  |  |  |  | 3 |  |  |  |  |  |  |  |  | 3 |
| China |  |  |  | 2 | 5 |  |  | 34 |  | 1 | 11 | 1 | 1 | 55 |
| Ecuador |  |  |  |  | 1 |  |  |  |  |  |  |  |  | 1 |
| Finland |  |  |  |  | 2 |  |  |  |  |  |  |  |  | 2 |
| France |  | 1 |  |  |  |  |  |  |  |  |  |  |  | 1 |
| Germany |  |  |  |  |  |  |  |  | 3 |  |  |  |  | 3 |
| India |  |  |  |  | 1 |  |  |  |  |  | 2 |  |  | 3 |
| Iraq |  |  |  |  | 1 |  |  |  |  |  |  |  |  | 1 |
| Mexico |  |  |  |  |  | 1 |  |  |  |  |  |  |  | 1 |
| Netherlands |  | 1 |  |  | 2 |  |  |  |  |  |  |  |  | 3 |
| Peru |  |  |  |  | 5 |  |  |  |  |  |  |  |  | 5 |
| Poland |  |  |  |  | 19 |  |  |  | 2 |  |  |  |  | 21 |
| Russia |  |  |  |  |  |  |  |  | 1 |  |  |  |  | 1 |
| Spain |  |  |  |  | 2 |  |  |  |  |  |  |  |  | 2 |
| Sweden |  |  |  |  | 1 |  |  |  |  |  |  |  |  | 1 |
| Thailand |  |  |  |  | 3 |  |  |  |  |  |  |  |  | 3 |
| United Kingdom |  |  |  | 1 | 3 |  |  |  | 1 |  |  |  |  | 5 |
| United States | 5 |  | 1 | 1 | 110 |  |  |  |  |  |  |  |  | 117 |
| **Total** | **6** | **3** | **1** | **4** | **173** | **1** | **3** | **34** | **7** | **1** | **13** | **1** | **1** | **248** |

**Table 2.** Summary of confirmed zoonotic IAVs and zoonotic classification category from literature review.

| Zoonotic Category | H1N1 | H1N2 | H3N2 | H3N8 | H5N1 | H5N2 | H5N5 | H5N6 | H5N8 | H7N9 | H9N2 | H10N5 | H10N8 | Total |
| --- | --- | --- | --- | --- | --- | --- | --- | --- | --- | --- | --- | --- | --- | --- |
| Avian-to-human |  |  |  | 3 | 33 | 1 |  | 33 | 1 | 1 | 13 | 1 | 1 | 86 |
| Avian-to-mammal |  |  |  | 1 | 114 |  | 3 | 1 | 6 |  |  |  |  | 125 |
| Mammal-to-human | 6 | 3 | 1 |  | 26 |  |  |  |  |  |  |  |  | 37 |
| **Total** | **6** | **3** | **1** | **4** | **173** | **1** | **3** | **34** | **7** | **1** | **13** | **1** | **1** | **248** |

We combined the confirmed zoonotic IAVs from our search with those in Eng *et al.* [17], for a total of 405 strains. The complete list and corresponding metadata of zoonotic strains can be found in the Supporting Information (SI) S1 Dataset and S1 Text, and a list of strains used in the zoonotic prediction model can be found in S2 Dataset.

### Influenza isolate risk prediction model

The overall structure of the influenza isolate risk prediction model is shown in Fig 1. Briefly, each influenza strain was separated into individual protein sequences and processed through ESM-2, a protein language model that was trained on protein sequences to predict protein structure, function, and effects of mutations [21]. The resulting embeddings were used to predict host tropism for each protein, which were then ensembled for the next model. The protein host tropism prediction values served as inputs for the zoonotic risk prediction model that classified strains into one of three host family groups or one of three zoonotic spillover categories, with confirmed zoonotic strains treated as a distinct classification. The model outputs the probability of each classification, providing the influenza isolate risk prediction value. Further details of each component are provided in the following sections.

**Fig 1.**
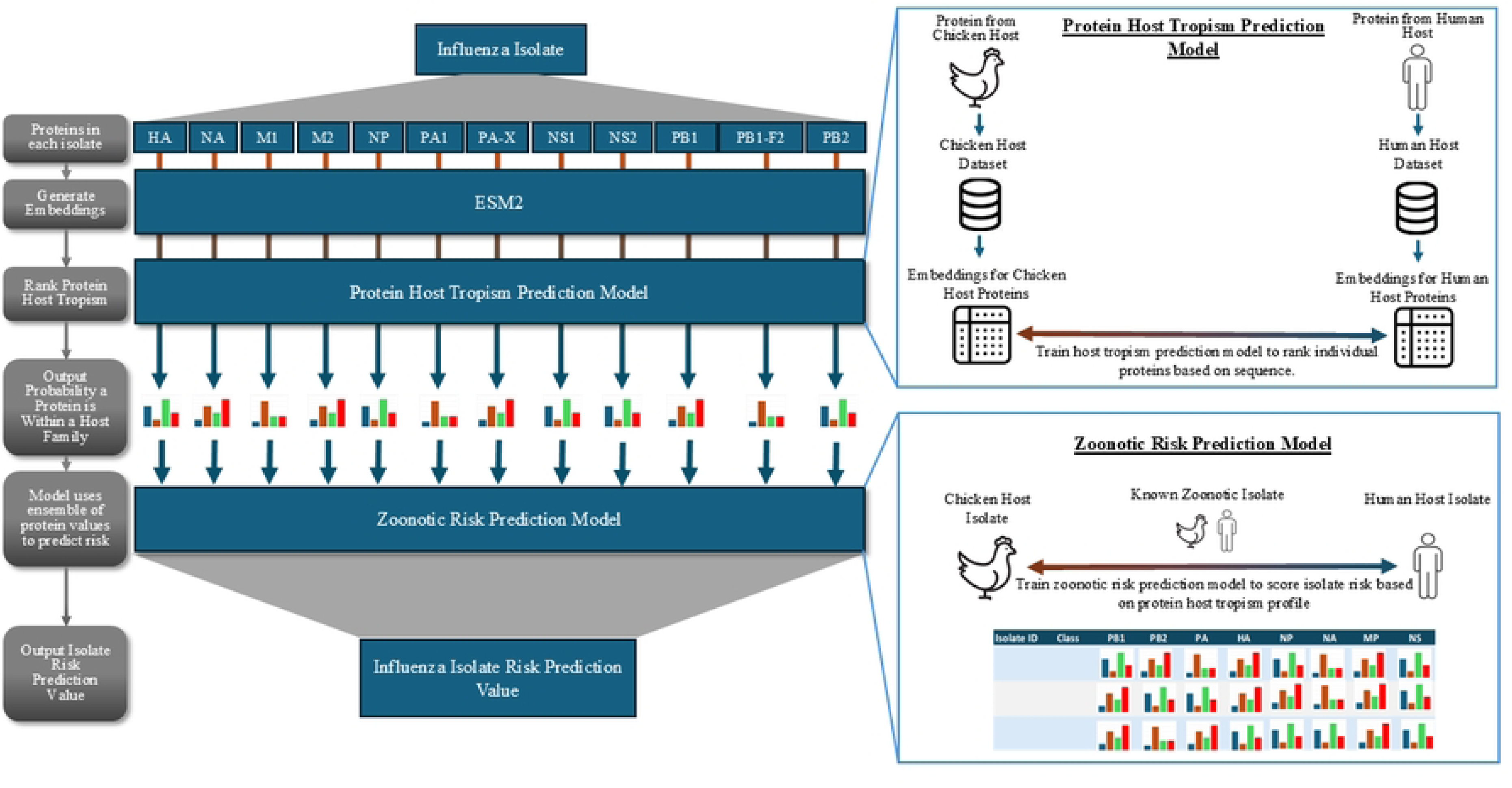
Influenza isolate risk prediction model workflow. The Protein Host Tropism Prediction Model inset illustrates the training approach using two host families as an example. The actual model was trained on nine host families (*Anatidae, Bovidae, Canidae, Equidae, Felidae, Hominidae, Laridae, Phasianidae, Suidae)*. The Zoonotic Risk Prediction Model inset similarly shows a simplified example, in which the actual model classifies strains into three host family groups (avian, mammal, and human) and three zoonotic spillover categories (avian-to-human, avian-to-mammal, and mammal-to-human) were used. The colored bars notionally represent the probability distribution generated by both models for each respective classification.

### Protein host tropism prediction model

A protein host tropism prediction model was developed to predict the source host of influenza strains by individual protein sequences. ESM-2 [21] was used to generate embeddings for the following 12 individual proteins in an isolate strain: hemagglutinin (HA), matrix proteins 1 (M1) and 2 (M2), neuraminidase (NA), nucleoprotein (NP), non-structural proteins 1 (NS1) and 2 (NS2), polymerase acidic protein (PA), polymerase basic proteins 1 (PB1) and 2 (PB2), PB1-F2, and PA-X. These embeddings served as input for eXtreme Gradient Boosting (XGBoost), a gradient-boosted decision tree ML algorithm. For each protein, the model generated output probabilities across nine taxonomic families representing the likelihood that the protein originated from each of the following host families: *Anatidae* (ducks and geese), *Bovidae* (domestic cattle), *Canidae* (dogs), *Equidae* (horses), *Felidae* (cats), *Hominidae* (humans), *Laridae* (gulls), *Phasianidae* (chickens and turkeys), and *Suidae* (domestic pigs). These families were selected based on their high spillover risk due to frequent contact with humans and recent outbreaks [2,18]. The probabilities outputs of this model were concatenated and served as input for the zoonotic risk prediction model.

### Zoonotic risk prediction model

A zoonotic risk prediction model was developed to evaluate influenza strain spillover risk using an XGBoost classifier. The model used outputs from the protein host tropism prediction model as input features to generate probabilities that a strain belongs to one of six classifications: avian, mammal, human, avian-to-human, avian-to-mammal, or mammal-to-human. The first three represent host family groups, while the latter three represent zoonotic spillover categories.

Host families were aggregated into broader groups due to limited availability of confirmed zoonotic strains for model training, as biosurveillance data remains sparse for certain species. The nine families were grouped as follows: avian (*Anatidae, Laridae, Phasianidae*), mammal (*Bovidae, Canidae, Equidae, Felidae, Suidae*), and human (*Hominidae*). Host family groups were selected from strains with at least 10 of the 12 proteins present (HA, M1, M2, NA, NP, NS1, NS2, PA, PB1, and PB2), as these were consistently present among sequences. The protein probabilities across all proteins were concatenated for input in the zoonotic model, providing at least 90 protein probabilities for each host family group strain. To prevent an imbalanced dataset, host family group strains were randomly downselected, while maintaining distributions of influenza subtypes and host families consistent with the overall dataset. As shown in Table 1, confirmed zoonotic strains are substantially less abundant than host-specific strains, as many spillover events are undetected, misdiagnosed, or unreported [22]. Due to this limited availability of confirmed zoonotic cases and limited sequencing of these strains, we could not limit zoonotic spillover strains to those that have at least 10 proteins. However, XGBoost manages missing data by determining the most effective default path for these values at each decision point, thereby maximizing predictive performance. An 80:20 random partition was applied to create the training and test datasets [23]. To assess model stability, 10 iterations were performed with different random partitions to ensure performance was not dependent on specific training data composition.

### Models and statistics analysis

All modeling and statistical analyses were conducted in Python version 3.11.7. Code can be accessed through GitHub at https://github.com/Noblis/influenza-spillover-prediction. All figures were created using Python and R Studio® version 2026.01.0+392. Embeddings from ESM-2 were generated with code available at https://github.com/facebookresearch/esm.

## Results

### Protein host tropism prediction model

To predict the host origin of influenza strains, we developed a protein host tropism prediction model and trained it on individual protein sequences from strains collected across nine taxonomic families. Table 3 shows the performance metrics for the model across all nine families. The model achieved high accuracy in classifying individual protein host tropism. A corresponding confusion matrix showing true vs. predicted labels is provided in the SI (S1 Fig). Throughout this analysis, we use the term “misclassified” to indicate instances where the predicted label does not match the true label. This terminology reflects prediction accuracy rather than underlying genetic relationships between host families. The model demonstrated strong predictive performance, with weighted precision and recall scores of 0.95 and 0.95, respectively. These metrics indicate that the model reliably predicted the likelihood of protein origin from each host family.

**Table 3.** Summary of protein host tropism model performance metrics.

| Host Family | Common name(s) | Total protein inputs | Precision | Recall | F1-score | Support |
| --- | --- | --- | --- | --- | --- | --- |
| *Anatidae* | Duck, goose | 212506 | 0.80 | 0.86 | 0.83 | 42501 |
| *Bovidae* | Domestic cow | 25168 | 0.86 | 0.99 | 0.92 | 5034 |
| *Canidae* | Dog | 5802 | 0.94 | 0.81 | 0.87 | 1160 |
| *Equidae* | Horse | 3680 | 0.95 | 0.93 | 0.94 | 736 |
| *Felidae* | Cat | 1640 | 0.91 | 0.13 | 0.23 | 328 |
| *Hominidae* | Human | 2108253 | 0.99 | 0.99 | 0.99 | 421651 |
| *Laridae* | Gull | 16500 | 0.90 | 0.63 | 0.74 | 3300 |
| *Phasianidae* | Chicken, turkey | 158475 | 0.73 | 0.72 | 0.72 | 31695 |
| *Suidae* | Domestic pig | 179756 | 0.94 | 0.87 | 0.90 | 35951 |
|  |  | **Accuracy** | 0.95 |  |  | 542356 |
|  |  | **Macro Average** | 0.89 | 0.77 | 0.79 | 542356 |
|  |  | **Weighted Average** | 0.95 | 0.95 | 0.95 | 542356 |

Precision scores, indicating the proportion of correct positive predictions, exceeded 0.80 for seven of nine families. *Anatidae* and *Phasianidae*, both common poultry, exhibited the lowest precision scores of 0.80 and 0.73, respectively; the third avian family, *Laridae*, achieved a higher precision of 0.90. *Hominidae*, or humans, had the highest precision, recall, and F1-scores among all families.

Recall scores, reflecting the model’s sensitivity in identifying true positive cases, exceeded 0.70 for seven of nine families. *Laridae* and *Felidae* exhibited the lowest recall scores of 0.63 and 0.13, respectively. The low recall for *Felidae* indicates that while positive predictions for this family were accurate (precision = 0.91; Table 3), the model misclassified 87% of cat-origin proteins as originating from other families. Notably, *Equidae* achieved high performance (precision = 0.95, recall = 0.93; Table 3) despite having only 3680 samples. Overall, the model performed well, with greater than 80% accuracy for six of the nine families and greater than 60% accuracy for two other families (S1 Fig).

Fig 2 shows the predicted protein host tropism for the eight primary proteins (HA, M, NA, NP, NS, PA, PB1, and PB2) derived from *Felidae*-associated strains, with color indicating the predicted host based on the highest probability score. *Felidae* was chosen because of its unique infection and transmission dynamics, as discussed later. Due to computational and graphical constraints associated with the large dataset, analysis was restricted to 100 randomly selected isolates possessing complete sequences for all eight proteins. Strain identifiers and corresponding host tropism for each protein are provided in S1 Table. As shown in Fig 2A, the majority of proteins were predicted to originate from *Bovidae* hosts, with only 5 stains exhibiting consistent *Felidae* prediction across all eight proteins. Numerous strains displayed heterogeneous host predictions across proteins, without discernible patterns. Fig 2B presents the probability distributions of six randomly selected isolates for each protein across all nine families. Strains 2, 3, and 6 showed high prediction probabilities for a single host family (*Felidae* or *Bovidae*), consistent with the observed high precision but low recall scores. Conversely, strains 1, 4, and 5 demonstrated lower single-family probabilities with greater distribution across the nine families, as indicated by lighter grey shading. Protein host tropism predictions and strain lists for the remaining eight host families are provided in S2-9 Figs and S2-9 Tables.

**Fig 2.**
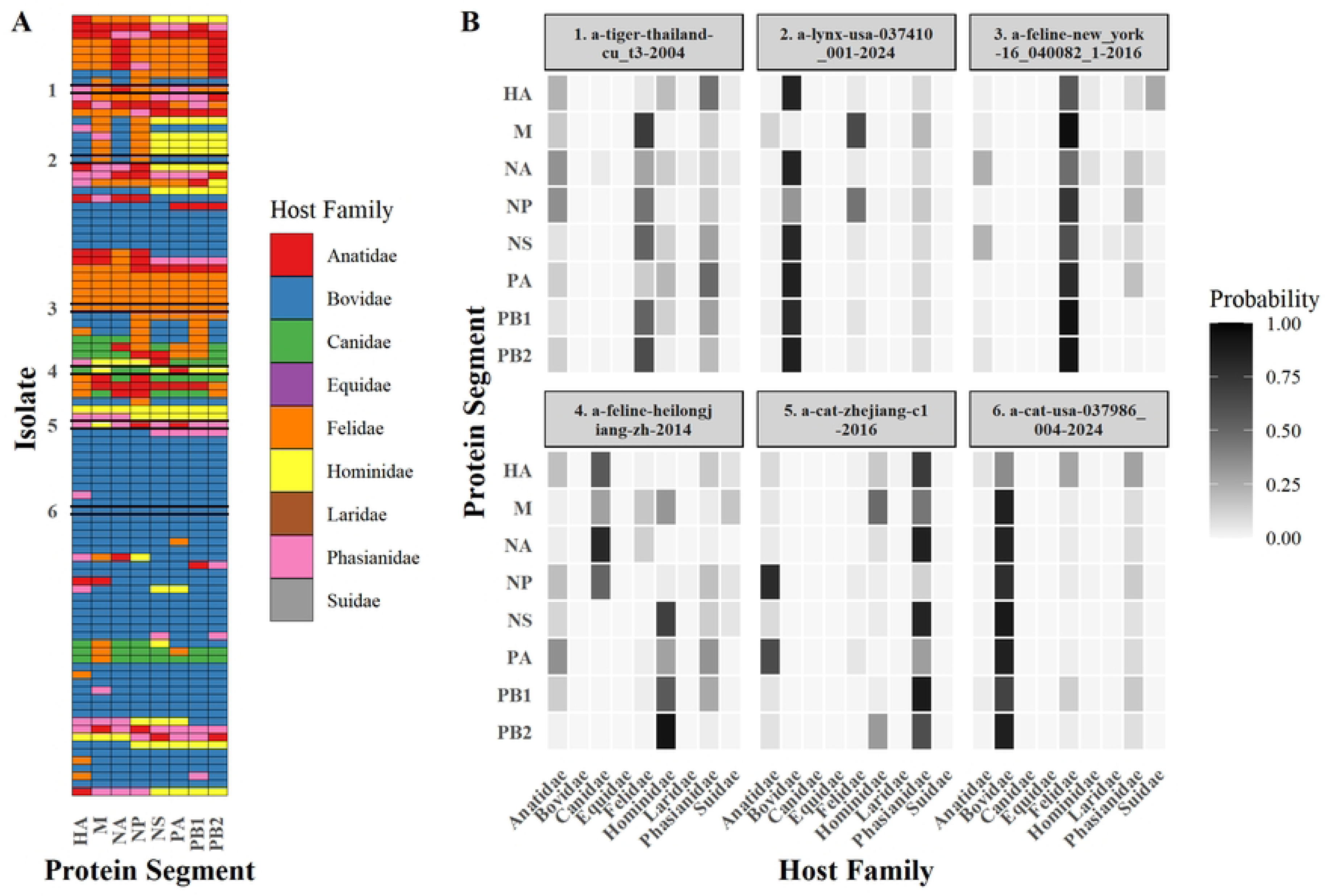
*Felidae* protein host tropism prediction results. (A) 100 strains containing all eight primary proteins (HA, M, NA, NP, NS, PA, PB1, and PB2) and (B) protein probability distributions for the eight primary proteins of six randomly selected isolates. For the left, the host tropism family is color-coded, and the highest probability host family is assigned to each protein. One row represents a single isolate. On the right, the individual protein probability distributions are shown for six isolates in the left figure. Cell colors represent the probability of belonging to each host family, ranging from light grey (lowest) to medium grey (intermediate) to black (highest). Isolate names are provided in S1 Table.

The remaining eight families demonstrated superior protein host tropism prediction accuracy compared to *Felidae* (13.1%)*. Anatidae* (S2 Fig) exhibited high classification accuracy (86.5%) with elevated host family prediction values; misclassified proteins were predominately assigned to *Phasianidae* or displayed low probabilities towards this family*. Bovidae* (S3 Fig) displayed high host family probabilities (98.9% accuracy), with only a single misclassified protein showing distributed probability between *Bovidae* and *Phasianidae. Canidae* (S4 Fig) demonstrated greater variability in protein host family probability while maintaining high prediction accuracy (80.5%), characterized by elevated protein host probabilities for *Canidae*. Both *Equidae* (S5 Fig) and *Hominidae* (S6 Fig) exhibited accurate host tropism predictions (92.9% and 98.9%, respectively) with consistently high protein prediction probabilities for correct classifications. *Laridae* (S7 Fig) demonstrated comparatively lower predictive performance (63.2%), with probability distributions frequently divided among other avian families, specifically *Anatidae* and *Phasianidae*. *Phasianidae* (S8 Fig) exhibited heterogeneous host tropism predictions (72.2% accuracy), with numerous proteins classified as *Anatidae*. Notably, even though *Suidae* showed high accuracy (87.0%), the majority of classifications outside of the host family were assigned to *Hominidae* (S9 Fig).

When protein predictions were aggregated into broader groups, avian, mammal, and human, model accuracy increased to 0.98. Precision values for the avian, mammal, and human groups were 0.95, 0.97, and 0.99, respectively, with results and performance metrics provided in the SI (S10-11 Tables). The model achieved strong overall predictive performance for protein host tropism, though performance varied across the nine animal families. These host tropism predictions were subsequently used as input features for assessing zoonotic spillover risk.

### Zoonotic risk prediction model

Using the ensembled protein host tropism probability outputs, the zoonotic risk prediction model classified strains into six classifications: avian, mammal, human, avian-to-human, avian-to-mammal, and mammal-to-human. The latter three categories represent the zoonotic spillover groups, defined as host range extension from one family to another. Through all 10 iterations, the model performed well with an average accuracy, weighted precision, and weighted recall of 0.90, 0.90, and 0.90, respectively (Table 4). Fig 3 shows a comparison between true labels and predicted labels generated by the model in iteration 8 out of 10 total, which achieved the highest performance.

**Fig 3.**
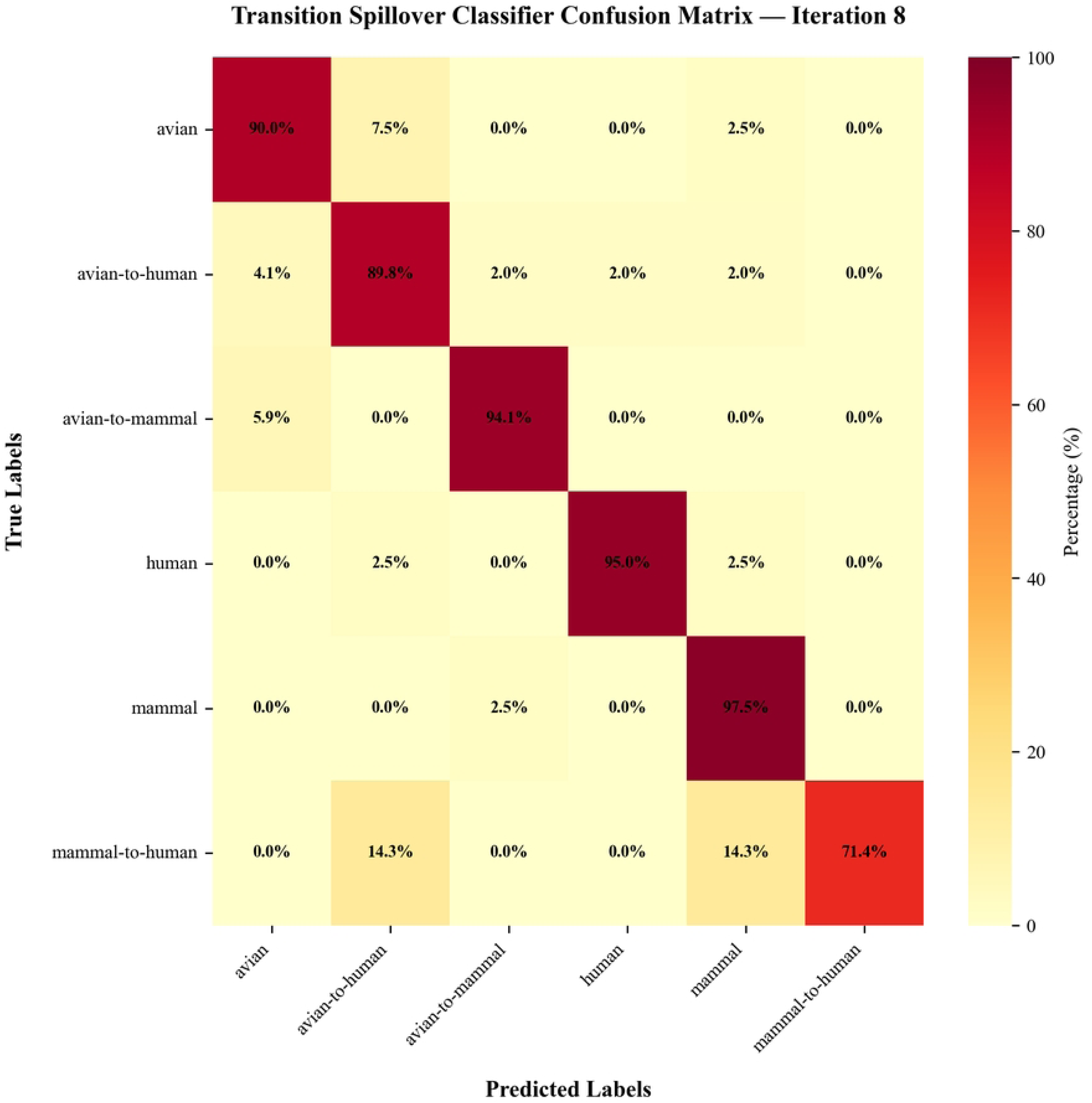
Percentage confusion matrix for zoonotic risk prediction model. Results shown are from iteration 8 of the spillover dataset. Rows represent true labels and columns represent predicted labels. Diagonal elements indicate correct classifications, while off-diagonal elements represent misclassifications. The test dataset is a random 20% selection of each host family group and zoonotic category in the spillover dataset (S2 Dataset; S12 Table).

**Table 4.**
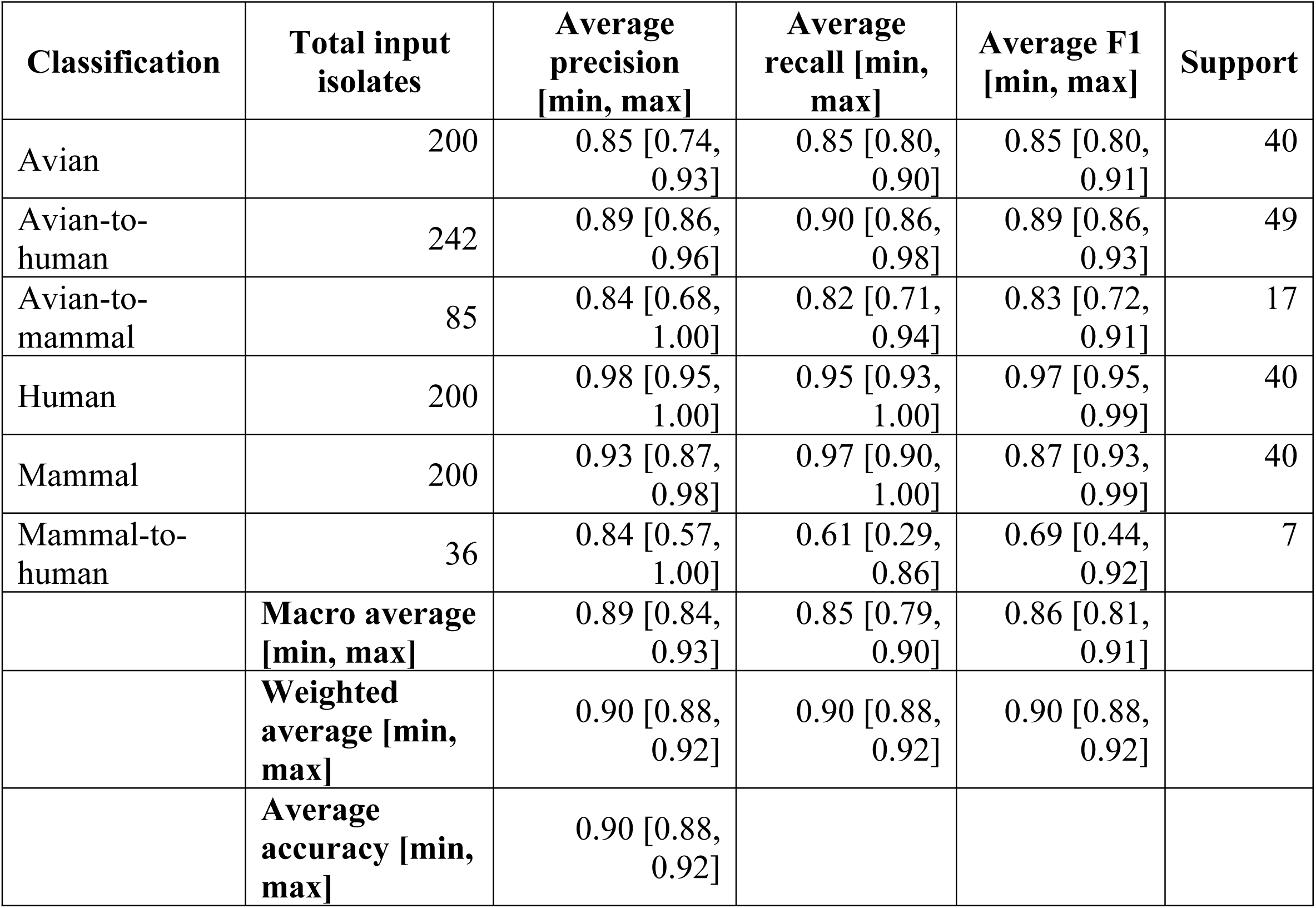
Summary of results for 10 iterations of zoonotic risk prediction model.

The human group achieved the highest performance metrics with average precisions and recall of 0.98 and 0.95 (Table 4), respectively, consistent with the individual protein host tropism predictive model results. The avian group demonstrated precision and recall of 0.85 and 0.85, respectively. The model correctly classified over 89% of the isolates in these spillover groups, as shown in the upper left quadrant of Fig 3. The mammal-to-human group exhibited distinct classification patterns, achieving maximum precision of 1.00 in iterations 1 and 8, with average precision and recall of 0.84 and 0.61, respectively, indicating high specificity but reduced sensitivity. The avian-to-mammal group showed average precision and recall of 0.84 and 0.82, respectively. To evaluate model performance on host family group classification, the model was run excluding spillover categories (S12 Table). Precision and recall for the avian and mammal groups without spillover categories were 0.98, 0.98, 0.97, and 0.97, compared to 0.85, 0.85, 0.93, and 0.97 when spillover groups were included. The inclusion of spillover groups reduced the classification performance for host-only groups, reflecting the increased complexity associated with cross-species transmission events.

Fig 4 shows the probability distribution for each test strain in iteration 8, the highest performing run with accuracy, precision, and recall scores of 0.92, 0.92, and 0.92, respectively. For strains in the avian, avian-to-human, and avian-to-mammal groups (Figs. 4A-C, S13 Table), the model predominately classified strains within related categories (i.e., an avian-to-human strain as avian, avian-to-human, or human), with few exceptions. S14 Table shows percentage confusion matrix for each classification across the 10 iterations. Strains in the human and mammal groups (Fig. 4D-E, S14 Table) were predicted with high confidence scores for their respective categories. The mammal-to-human groups exhibited strong predictions for either the avian-to-human spillover (24.3%) or mammal (8.6%) classifications. Avian spillover groups consistently exhibited higher prediction probabilities towards their original host group compared to their current host group (avian-to-human as avian 6.9%, avian-to-mammal as avian 12.9%). Overall, the majority of strains across all groups demonstrated high probability predictions, indicating robust model confidence in group classification.

**Fig 4.**
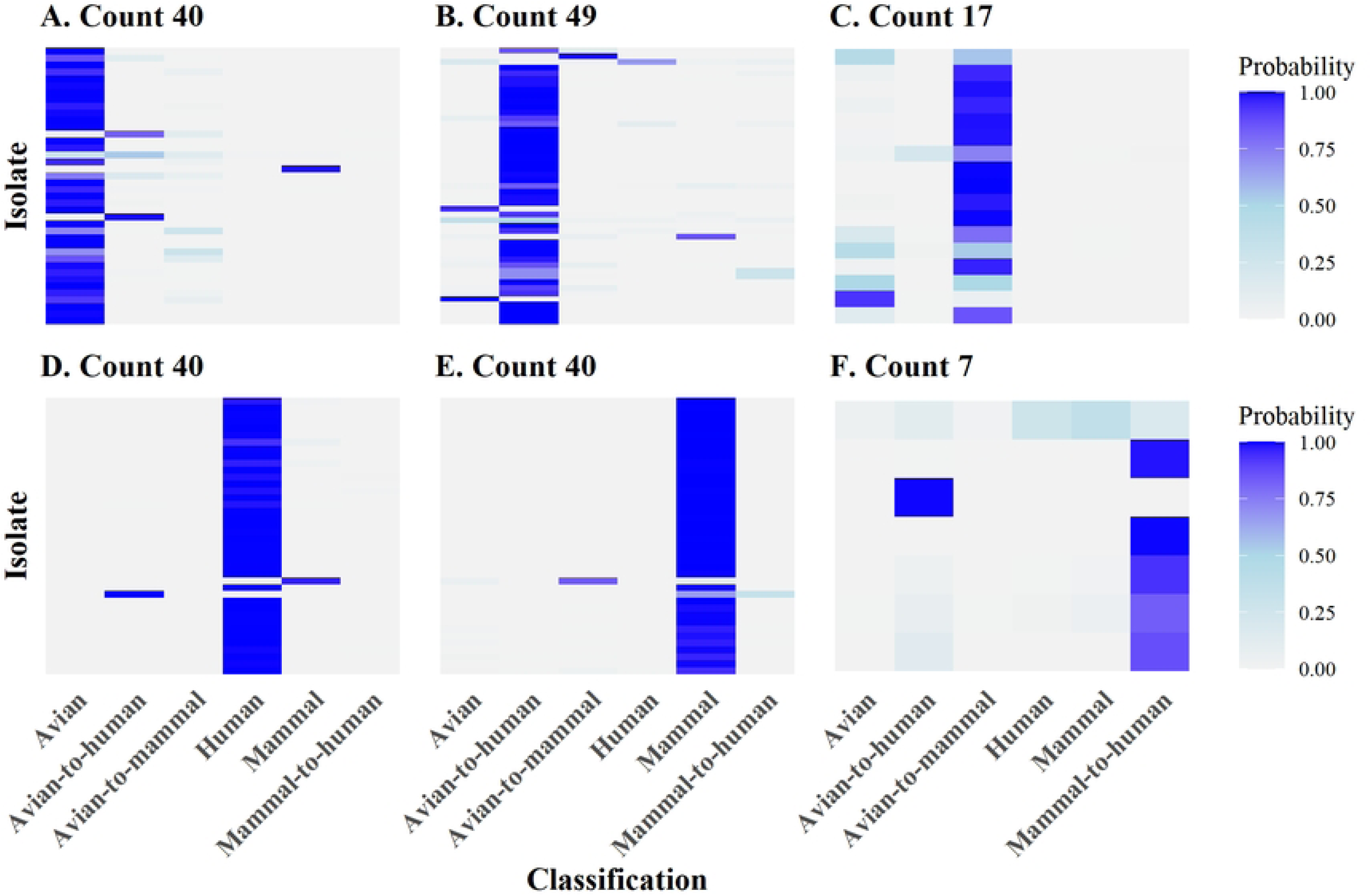
Zoonotic risk prediction model strain classification probability output iteration 8 of 10. (A) Avian, (B) avian-to-human, (C) avian-to-mammal, (D) human, (E) mammal, and (F) mammal-to-human spillover groups probability outputs along with the total number of strains used in model. The color range represents probability from low (light grey) to high (medium blue). Strains and probability values can be found in S13 Table. The dataset only includes test results and was the highest performing iteration.

A total of 101 isolate strains were misclassified (i.e., classified as associated with a host other than observed) in at least one iteration of the model. S15 Table summarizes the strains demonstrating classification errors and their corresponding misclassification frequencies across 10 iterations. Twenty-five strains (24.8%) exhibited misclassification rates ≥30%, with three strains demonstrated a 50% misclassification rate. All misclassifications were confined to test set predictions, with 79 strains (∼78%, highlighted in orange) being consistently misclassified across all iterations in which they appeared in the test sets.

## Discussion

### Model analytics

We developed two predictive models to assess influenza spillover risk, a critical component of safeguarding economic and public health systems against emerging infectious disease threats. The first model generated individual protein host tropism prediction profiles from generated ESM-2 embeddings of proteins. The second model ensembled and integrated the protein host tropism profiles with confirmed zoonotic strain data to classify strains into host family groups or zoonotic spillover categories.

A key advance of our approach was training the tropism model on all available subtypes, variants, and mutations of influenza virus across the nine host families. This enables the model to abstract tropism profiles from the broadest possible dataset. Protein structure is a critical determinant of viral fitness and host range [24,25], yet the precise mechanisms underlying spillover remain poorly understood. Influenza spillover may result from surface proteins acquiring affinity for receptors in new hosts, changes to the polymerase, subtle structural changes affecting protein function, or a combination of all [2]. We hypothesized that ESM-2, a model that more closely represents the protein structure compared with genomic models, protein one-hot encoding, or similar approaches, would provide better predictive performance across diverse viral lineages and potentially lead to a model that is agnostic to virus type. ESM-2’s training enables accurate prediction of the impact of mutations absent from its training data [26]. Additionally, as a protein language model, ESM-2 offers lower computations cost than other protein models, allowing for scalability [21].

We selected XGBoost as our classification algorithm due to its distinct advantages in handling large, complex datasets with high predictive accuracy and computational efficiency [27]. These characteristics were particularly well-suited for our protein host tropism model, which processed nearly 3 million proteins across a high-dimensional representation space. XGBoost also demonstrates superior performances in handling imbalanced datasets compared to other ML architectures [28], making it appropriate for both of our prediction models where class imbalance was present. This choice is further supported by Eng *et al.* [29], who compared random forest, another decision tree ensemble method, against other ML classifiers (Naïve Bayes, kNN, SVM, and ANN), finding random forest outcompeted all the other classifiers. Given the established performance advantages of tree-based ensemble methods in similar biological prediction tasks, XGBoost represented an optimal choice for our framework and exhibited high predictive performances.

Our model demonstrated comparable accuracy to previously developed deep-learning based protein prediction models [14,15,17,29,30]. Hu *et al.* [14] developed Flu-CNN, which achieved 99% accuracy in determining host specificity from a single genomic segment and identified mosaic patterns in the strains that highlight the potential zoonotic risk. Similarly, Eng *et al.* [29] developed computational models for 11 influenza proteins to predict host tropism with 99% accuracy, later extending this work to predicting zoonotic risk with 97% accuracy [17]. However, these models were limited to binary avian-human classification, excluding mammals. As will be discussed later, mammalian hosts often serve as intermediate species between birds and humans. These intermediate hosts not only pose significant spillover risk but also introduce additional complexity that challenges prediction modeling [22], potentially explaining the higher performance of Eng’s binary zoonotic classification compared to our multi-host approach. When proteins were classified into broad groups similar to Eng’s approach, our protein prediction model achieved comparable accuracy (98%). Furthermore, avian-to-human spillover cases are more frequently documented due to the rise in bird flu [31], providing larger training datasets for models solely focused on this transmission pathway.

Nguyen *et al.*’s WaveSeekerNet incorporated mammals and achieved 97% accuracy using only HA and NA proteins for host prediction [15]. While not explicitly designed for zoonotic spillover prediction, further analysis revealed that host prediction discrepancies often corresponded to recent zoonotic transmission events [15]. Although informative, such retrospective analysis lacks the rapid response required for proactive zoonotic surveillance and risk interventions. Like previous models [14,17], WaveSeekerNet employed broad classification groups, limiting the host specificity and associated spillover risk. Alberts *et al.* [30] developed a gradient boosting machine classifier with 98% accuracy across seven host families (canine, chicken, equine, human, mallard, swine, and turkey) but restricted analysis to the HA protein of H3 subtype strains. This provided a narrow dataset given the diversity of circulating subtypes and missing those that potentially pose greater pandemic risk (e.g., H5N1, H7N9). This model also did not attempt zoonotic risk prediction.

Our zoonotic risk model addressed these limitations by incorporating confirmed spillover strains as distinct training categories, grounding predictions in empirical data spanning reported influenza subtypes worldwide and providing a wider range of host specificity. Critically, we included mammalian spillover events alongside avian-to-human transmission, expanding both dataset size and risk scope. Mammalian hosts represent critical intermediate species between avian reservoirs and humans [18,32], providing environments where viruses can acquire mammalian adaptations, undergo reassortment with other subtypes, or accumulate mutations that enhance human transmissibility, as exemplified by the 2009 swine flu pandemic [33,34]. While the finer-scale resolution of host families and addition of mammalian hosts decreased overall accuracy compared to other models, both models maintained high accuracy (≥0.90) while enabling more quantitative risk assessment.

The model generates probabilistic classifications across host family groups and zoonotic categories for each strain, with the highest probability indicating the predicted risk level. With additional work, these quantitative risk values could be used to establish risk thresholds to inform further laboratory analysis and proactive intervention strategies. Identifying spillover events is a crucial task in preventing potential outbreaks in naïve populations [2]. Overall, our framework offers computational and economic advantages over existing models while providing more granular host-specific predictions. The following sections examine each individual model’s performance in detail.

### Performance of protein host tropism prediction model

To our knowledge, this is the first model to predict influenza strain host tropism across nine animal families with high accuracy. Identifying strain host origin is important not only for quantifying risk within that species, but also for assessing zoonotic potential, as certain animal hosts pose greater spillover risk to humans or other species [35].

Our approach employed a hierarchical framework, beginning with protein-level host tropism predictions. Influenza viruses and their proteins undergo molecular adaptations, such as mutation and reassortment, and evolve in a host as they circulate within the population [36,37]. As a result, proteins may be most closely aligned to a single host, or in the case of a spillover event, a mixture of the current and previous host, depending on the time of spillover and the rate of viral evolution, further complicating strain host-origin predictions. By first identifying protein-level patterns and mutations, our model captures evolutionary heterogeneity across viruses and generates more holistic prediction profiles. This resolution may enable future applications in predicting spillover risk between closely related families, such as *Anatidae, Laridae,* and *Phasianidae*.

Most existing prediction models treat avian hosts as a single group, obscuring important variations among species that may influence viral ecology and transmission dynamics. By resolving avian hosts into three distinct families, our model captures finer-scale differences that inform a more comprehensive and informative cross-species risk assessment. *Anatidae* (ducks and geese) and *Laridae* (gulls) are natural reservoirs for influenza A viruses (IAV) [37], yet direct spillover from wild aquatic birds to humans are less common due to limited contact and insufficient viral adaptations for human infection [2]. While strains adapted to *Phasianidae* (turkeys and chickens) also lack the required viral adaptations, they elicit a greater threat due to increased contact through domestic poultry production and proximity to mammalian livestock [38]. Moreover, outbreaks within poultry populations can lead to major economic loss [3], making rapid identification crucial for both public health and agricultural biosecurity.

The model achieved high precision and recall across avian hosts, demonstrating strong performance in distinguishing among avian families. Given the conserved receptor-binding preference and other shared adaptations among strains adapted to avian species [2], we anticipated that some avian strains would still exhibit low-probability assignments to other avian families. As shown in S1 Fig, *Anatidae* proteins were primarily misclassified as *Phasianidae* (and vice versa), while *Laridae* proteins were misclassified as *Anatidae* and *Phasianidae*. *Anatidae* strains (S2 Fig) with lower probabilities for their canonical host often exhibited secondary probabilities for *Phasianidae,* with minimal assignment to *Laridae.* In contrast, *Phasianidae* strains (S8 Fig) displayed broader probability distributions across multiple families—predominately *Anatidae,* but also *Hominidae, Laridae,* and *Suidae*—while *Laridae* strains (S7 Fig) showed probability profiles largely restricted to *Anatidae* and *Phasianidae.* The finer-scale family host resolution of our model raises the question of whether these represent true protein misclassifications or whether certain proteins are more representative of alternative hosts, a consideration inherent to using specific host families rather than broad taxonomic groups. Further work is needed to elucidate these distinctions.

These patterns do align with expected genetic similarity between families, spillover dynamics, and ecological relationships among host species. As primary natural reservoirs [37], strains adapted to ducks and geese appear to possess distinctive protein-level signatures, with misclassifications occurring primarily with strains adapted to turkeys and chickens, species with which they may frequently interact in agricultural settings. In keeping with leading theories on AIV evolution and spread, the broader probability distributions observed for poultry may reflect their increased interspecies contact on farms [38], elevating spillover risk. No discernible pattern in protein misclassification was observed, indicating that classification errors were not attributed to single protein species-specific adaptive signatures. Additionally, diffuse probability profiles may indicate emerging host adaptations preceding spillover events or recently completed host jumps. Although formal zoonotic risk assessment occurs in the subsequent model, these host-tropism profiles may serve as an early indicator of strains with heightened spillover potential. Given that viral sampling, sequencing, and analysis require time during which transmission may already be occurring in naïve populations, leveraging protein prediction profiles for proactive threat identification could enable earlier and more targeted interventions. Further work is needed to quantify how these probabilistic prediction profiles translate to measurable spillover risk metrics.

Domestic mammalian families exhibited higher protein prediction accuracy (>80% except *Felidae*, S1 Fig) and more constrained host tropism profiles compared to avian families. *Equidae* (S5 Fig) and *Bovidae* (S3 Fig) showed strong source-host probabilities, with *Bovidae* displaying one protein prediction and low probability distribution for *Phasianidae*, indicating chickens and turkeys—not wild birds as hypothesized [39]—could be spillover sources in agricultural environments. Further investigation and analysis are needed to confirm, as these results could have significant biosecurity implications. Interestingly, *Suidae* (S9 Fig) exhibited high source-host probability and prediction but secondary signal and prediction for *Hominidae,* with *Hominidae* as the most frequent misclassification (11.9%) for *Suidae* proteins (S1 Fig). Together, these patterns likely reflect the frequent bidirectional spillover between humans and pigs [40,41]. Probability assignments to avian families remained surprisingly low despite pigs expressing both avian- and mammalian-type sialic acid receptors [19], which enables infection from all host family groups. This pattern may reflect rapid post-spillover adaptation in swine hosts, though it is likely influenced by sampling bias, as many analyzed strains could have been collected after multiple rounds of host-specific adaptations had occurred. Future work incorporating larger, temporally resolved datasets will be essential in resolving these evolutionary and ecological dynamics.

Although *Canidae* had high precision and recall values in Table 3, this is not displayed in the first rows of S4 Fig. A subset (10%) of the total 100 *Canidae* samples in S4 Fig include non-domestic species such as foxes. Strains isolated from domestic dogs, highlighted in the majority of S4 Fig, exhibited high source-host probabilities and predictions consistent with overall performance metrics, with some notable exceptions. Foxes and other wild animals within the *Canidae* family typically acquire infections from consumption of infected birds [42,43], resulting in avian-to-mammal spillover events rather than sustained circulation within the population. This emphasizes the different transmission routes that can occur between different species within the same animal family, an important implication for risk assessment.

The notable exception in the host tropism model’s performance was *Felidae*, which had a low recall of 0.13. We propose two primary explanations for this limitation. First, regular testing of domestic cats for AIV has only recently occurred, after cat infections were identified to be a potential marker for the presence of the virus in a specific location [44,45]. As a result, training data for this family remained substantially smaller than for other hosts. However, as previously mentioned, *Equidae* exhibited high protein prediction accuracy (∼93%) despite having only 3680 samples, substantially fewer than most families yet only approximately twice that of *Felidae.* This differentiation may reflect varying infection and transmission dynamics between the species, as equine influenza is highly contagious and spreads rapidly among horses [46], providing more opportunities to observe host-specific adaptations.

Second, farm cats are hypothesized to be infected by ingesting raw milk or poultry [47–49]. While some studies have reported cat-to-cat transmission [49–51], they are limited in number. Combined with reduced surveillance efforts, this likely resulted in fewer observable cat-host adaptation patterns for model identification. Consistent with this interpretation, most *Felidae* proteins were classified as *Anatidae* (ducks and geese), *Phasianidae* (chickens and turkeys), or *Bovidae* (domestic cows) (Fig 2 and S1 Fig). Infections in domestic cats likely represent recent spillovers rather than endemic circulation within the *Felidae* family. This could explain why the host tropism prediction was not consistent within each *Felidae* species (Fig 2). Further work is needed to elucidate whether reduced predictive performance stems from small sample size, unique transmission dynamics, or both. We considered removing Felidae from the dataset; however, we retained it for dataset completeness and in anticipation of expanding surveillance efforts. Although *Felidae* currently poses a minimal spillover risk to humans, some cases of human infections linked to domestic cats have occurred [52–54]. This indicates the importance of continuous monitoring and use of *Felidae* in our model. Overall, the host tropism prediction profiles generated by this model provided valuable input for the zoonotic spillover risk prediction model.

### Performance of zoonotic risk prediction model

While identifying zoonotic spillover threats is crucial for risk mitigation, accurate prediction remains challenging. Our zoonotic risk prediction model performed well in distinguishing between zoonotic spillover and host family group strains. During early spillover events, strains typically retain genetic signatures of their original host, as insufficient time has elapsed for adaptation to their current host [55]. This pattern was evident in our results, with strains in the avian-to-human and avian-to-mammal categories exhibiting higher probability distributions toward their original host family groups. The reduced predictive performance for the avian host family group compared to the human and mammal host family groups can likely be attributed to avian strains serving as the original host for two spillover categories, compared to none and one for the human and mammal groups, respectively.

Notably, strains in the avian-to-human category showed elevated probabilities for avian or avian-to-mammal classifications with minimal probability or classification (∼0.6%) of the human group, further demonstrating the persistent influence of the original host characteristics on viral genetic features. Interestingly, the mammal-to-human group had higher prediction of avian-to-human (24.3%, highest misclassification in the model) compared to mammals (8.6%) or humans (4.3%). We hypothesize this is due to the limited data reducing prediction performance and errors in the input dataset, as discussed below, rather than similar genetic patterns between the categories. Importantly, while the model misclassified mammal-to-human spillover cases as avian-to-human, it correctly identified human infection risk. When these two spillover groups are combined, the lowest predictive accuracy increases from 71.4% to 85.7%, indicating that the model successfully flags strains with human spillover potential, the primary objective of this classification framework.

The avian-to-mammal and mammal-to-human spillover groups exhibited the poorest performance. Both groups contained the smallest number of strains, directly contributing to reduced model accuracy. To enhance model performance, continued literature review and incorporation of emerging spillover surveillance data are essential, as the power of ML models is driven by the dataset quality and size. Additionally, the zoonotic spillover model’s performance is constrained by the host tropism model’s predictions. While the host tropism model achieved overall high accuracy, certain host families, such as *Felidae,* demonstrated low recall that likely propagated errors into the zoonotic risk predictions. Expansion of the host tropism model dataset and refinement of its architecture are critical as biosurveillance capabilities advance.

Analysis of misclassified strains revealed important patterns (S15 Table). No discernable trend emerged, suggesting that specific host families were disproportionately associated with prediction errors. However, 79 strains were consistently misclassified across all test set iterations, with 52 of these misclassified in multiple iterations (S15 Table). These persistently misclassified strains warrant further investigation to identify the underlying genomic or epidemiological features that may limit model learning and to assess whether they represent errors in the input data from the protein host tropism model.

Our model employed a more comprehensive dataset than previous approaches by examining spillover events across multiple animal families and diverse zoonotic transmission pathways, rather than focusing exclusively on avian-to-human transmission [14,17]. These additional classification categories captured more complex interaction networks but necessitated a larger training dataset. However, we acknowledge that influenza viruses infect and spill over into animal species beyond the nine host families included in this study [35]. Expansion of datasets with greater depth sequencing depth across additional host families is necessary to evaluate model generalizability, though the predominance of wild species among underrepresented hosts present inherent challenges due to limited genomic surveillance in these populations. Given the zoonotic spillover model’s dependance on the host tropism predictions, we did not include zoonotic spillover cases outside the nine host families in the training data and hypothesized it would decrease the model’s performance. Therefore, we recommend expanding the host family representation in the foundational host tropism model rather than attempting predictions for novel host families post hoc.

The primary output of the zoonotic risk model is the influenza isolate risk prediction value, defined as the maximum probability among all zoonotic group classifications and indicated by darker shading in Fig 4. Elevated predictive values in spillover groups may indicate a strain’s potential for sustained circulation within a naïve host population. However, additional analyses are required to establish the relationship between this probability score and transmission potential within new host populations, ultimately enabling development of a combined spillover and transmission score risk that integrates both emergence likelihood and epidemic potential.

## Conclusion: real-world implications

Avian influenza viruses pose a substantial threat to public health, as each spillover event increases the probability of human exposure to infected animals and expands the diversity of influenza strains capable of zoonotic transmission. Through continued mutation and reassortment, AIVs may acquire human-to-human transmission capability, potentially triggering a pandemic. Therefore, early detection, comprehensive surveillance, and strategic resource allocation guided by predictive modeling are critically important for pandemic preparedness. Rapid identification and risk assessment of novel spillover strains is essential, as newly introduced influenza viruses can spread rapidly through immunologically naïve populations, resulting in significant harm to species’ health. Our novel modeling framework demonstrated robust performance in predicting both host tropism and zoonotic risk for influenza strains across nine host families and three spillover categories. Key takeaways from our models are: 1) host tropism prediction is likely influenced by the sample size as well as the infection route and transmission dynamics, 2) we are able to distinguish host prediction on a family level, which enhances risk prediction modeling, and 3) the addition of mammals in both models increased the complexity of the dataset and provided a more quantitative risk assessment. To enhance model utility, future work should prioritize the expansion of the zoonotic spillover dataset and extension of the host tropism model to include additional animal families that represent emerging spillover threats.

## Limitations

Dataset limitations, particularly the constrained sample sizes for certain zoonotic spillover groups and host families, represented a significant challenge in this work. These limitations necessitated reducing the number of input features for the zoonotic risk model, consequently affecting predictive performance. Future work should systematically evaluate the impact of dataset imbalance and class representation on the performance of both the protein host tropism and zoonotic risk models. Additionally, this study employed a single ML architecture for predictions. Comparative analyses of alternative gradient boosting algorithms, such as LightBGM, CatBoost, or random forest models, should be conducted for both the host tropism and zoonotic risk components of the modeling framework. These algorithms have different strengths and weaknesses with respect to nonlinearity, interactions, and dataset size. Several aspects of the model performance remain unknown, including the consistent misclassification of several strains across all iterations of the zoonotic model. Additionally, future work should conduct more comprehensive analyses, including phylogenetic investigations to assess whether predictive accuracy differs between viral lineages associated with multiple host species versus lineages restricted to single hosts. This could elucidate whether spillover events leave detectable genomic signatures that influence model predictions.

## Acknowledgements

We gratefully acknowledge the work of all authors who generated and submitted the numerous sequences to GISAID’s EpiFlu™ Database and the NCBI’s GenBank Protein Database on which this research was based on.

## Supporting information

**S1 Dataset. Metadata of confirmed zoonotic cases from literature review. S1 Text. Reference list for S1 Dataset.**

**S2 Dataset. List of strains used in zoonotic prediction risk model.**

**S1 Fig. Confusion matrix for protein host tropism model.** Rows represent true labels and columns represent predicted labels. The test dataset is a random 20% selection of individual proteins from each host family.

S1 Table. *Felidae* isolates protein host tropism predictions from Fig 2.

**S2 Fig. *Anatidae* protein host tropism prediction results.** (A) 100 strains containing all eight primary proteins (HA, M, NA, NP, NS, PA, PB1, and PB2) and (B) protein probability distributions for the eight primary proteins of six randomly selected isolates. For the left, the host tropism family is color-coded, and the highest probability host family is assigned to each protein. One row represents one isolate. On the right, the individual protein probability distributions are shown for six isolates in the left figure. Cell colors represent the probability of belonging to each host family, ranging from light grey (lowest) to medium grey (intermediate) to black (highest). Isolate names are provided in S2 Table.

S2 Table. *Anatidae* isolates protein host tropism predictions from S2 Fig.

**S3 Fig. *Bovidae* protein host tropism prediction results**. (A) 100 strains containing all eight primary proteins (HA, M, NA, NP, NS, PA, PB1, and PB2) and (B) protein probability distributions for the eight primary proteins of six randomly selected isolates. For the left, the host tropism family is color-coded, and the highest probability host family is assigned to each protein. One row represents one isolate. On the right, the individual protein probability distributions are shown for six isolates in the left figure. Cell colors represent the probability of belonging to each host family, ranging from light grey (lowest) to medium grey (intermediate) to black (highest). Isolate names are provided in S3 Table.

S3 Table. *Bovidae* isolates protein host tropism predictions from S3 Fig.

**S4 Fig. *Canidae* protein host tropism prediction results.** (A) 100 strains containing all eight primary proteins (HA, M, NA, NP, NS, PA, PB1, and PB2) and (B) protein probability distributions for the eight primary proteins of six randomly selected isolates. For the left, the host tropism family is color-coded, and the highest probability host family is assigned to each protein. One row represents one isolate. On the right, the individual protein probability distributions are shown for six isolates in the left figure. Cell colors represent the probability of belonging to each host family, ranging from light grey (lowest) to medium grey (intermediate) to black (highest). Isolate names are provided in S1 Table.

S4 Table. *Canidae* isolates protein host tropism predictions from S4 Fig.

**S5 Fig. *Equidae* protein host tropism prediction results.** (A) 100 strains containing all eight primary proteins (HA, M, NA, NP, NS, PA, PB1, and PB2) and (B) protein probability distributions for the eight primary proteins of six randomly selected isolates. For the left, the host tropism family is color-coded, and the highest probability host family is assigned to each protein. One row represents one isolate. On the right, the individual protein probability distributions are shown for six isolates in the left figure. Cell colors represent the probability of belonging to each host family, ranging from light grey (lowest) to medium grey (intermediate) to black (highest). Isolate names are provided in S5 Table.

S5 Table. *Equidae* isolates protein host tropism predictions from S5 Fig.

**S6 Fig. *Hominidae* protein host tropism prediction results.** (A) 100 strains containing all eight primary proteins (HA, M, NA, NP, NS, PA, PB1, and PB2) and (B) protein probability distributions for the eight primary proteins of six randomly selected isolates. For the left, the host tropism family is color-coded, and the highest probability host family is assigned to each protein. One row represents one isolate. On the right, the individual protein probability distributions are shown for six isolates in the left figure. Cell colors represent the probability of belonging to each host family, ranging from light grey (lowest) to medium grey (intermediate) to black (highest). Isolate names are provided in S6 Table.

S6 Table. *Hominidae* isolates protein host tropism predictions from S6 Fig.

**S7 Fig. *Laridae* protein host tropism prediction results.** (A) 100 strains containing all eight primary proteins (HA, M, NA, NP, NS, PA, PB1, and PB2) and (B) protein probability distributions for the eight primary proteins of six randomly selected isolates. For the left, the host tropism family is color-coded, and the highest probability host family is assigned to each protein. One row represents one isolate. On the right, the individual protein probability distributions are shown for six isolates in the left figure. Cell colors represent the probability of belonging to each host family, ranging from light grey (lowest) to medium grey (intermediate) to black (highest). Isolate names are provided in S7 Table.

S7 Table. *Laridae* isolates protein host tropism predictions from S7 Fig.

**S8 Fig. *Phasianidae* protein host tropism prediction results**. (A) 100 strains containing all eight primary proteins (HA, M, NA, NP, NS, PA, PB1, and PB2) and (B) protein probability distributions for the eight primary proteins of six randomly selected isolates. For the left, the host tropism family is color-coded, and the highest probability host family is assigned to each protein. One row represents one isolate. On the right, the individual protein probability distributions are shown for six isolates in the left figure. Cell colors represent the probability of belonging to each host family, ranging from light grey (lowest) to medium grey (intermediate) to black (highest). Isolate names are provided in S8 Table.

S8 Table. *Phasianidae* isolates protein host tropism predictions from S8 Fig.

**S9 Fig. *Suidae* protein host tropism prediction results.** (A) 100 strains containing all eight primary proteins (HA, M, NA, NP, NS, PA, PB1, and PB2) and (B) protein probability distributions for the eight primary proteins of six randomly selected isolates. For the left, the host tropism family is color-coded, and the highest probability host family is assigned to each protein. One row represents one isolate. On the right, the individual protein probability distributions are shown for six isolates in the left figure. Cell colors represent the probability of belonging to each host family, ranging from light grey (lowest) to medium grey (intermediate) to black (highest). Isolate names are provided in S9 Table.

S9 Table. *Suidae* isolates protein host tropism predictions from S9 Fig.

**S10 Table. Individual protein host tropism prediction model results for broader host family grouping.**

**S11 Table. Individual protein host tropism prediction model performance metrics for broader host family grouping.**

**S12 Table. Host family groups only zoonotic risk prediction model performance metrics for 10 iterations.**

**S13 Table. Summary of results from iteration 8 of 10 of zoonotic risk prediction model.**

**S14 Table. Zoonotic risk model combined classifications percentages for across 10 iterations.**

**S15 Table. Summary of misclassified virus strains in zoonotic risk prediction across 10 iterations.**

